# Gene birth contributes to structural disorder encoded by overlapping genes

**DOI:** 10.1101/229690

**Authors:** S. Willis, J. Masel

## Abstract

The same nucleotide sequence can encode two protein products in different reading frames. Overlapping gene regions encode higher levels of intrinsic structural disorder (ISD) than non-overlapping genes (39% vs. 25% in our viral dataset). This might be because of the intrinsic properties of the genetic code, because one member per pair was recently born de novo in a process that favors high ISD, or because high ISD relieves increased evolutionary constraint imposed by dual-coding. Here we quantify the relative contributions of these three alternative hypotheses. We estimate that the recency of de novo gene birth explains 32% or more of the elevation in ISD in overlapping regions of viral genes. While the two reading frames within a same-strand overlapping gene pair have markedly different ISD tendencies that must be controlled for, their effects cancel out to make no net contribution to ISD. The remaining elevation of ISD in the older members of overlapping gene pairs, presumed due to the need to alleviate evolutionary constraint, was already present prior to the origin of the overlap. Same-strand overlapping gene birth events can occur in two different frames, favoring high ISD either in the ancestral gene or in the novel gene; surprisingly, most de novo gene birth events contained completely within the body of an ancestral gene favor high ISD in the ancestral gene (23 phylogenetically independent events vs. 1). This can be explained by mutation bias favoring the frame with more start codons and fewer stop codons.

## I. Introduction

Protein-coding genes sometimes overlap, i.e. the same nucleotide sequence encodes different proteins in different reading frames. Most of the overlapping pairs of genes that have been characterized to date are found in viral, bacterial and mitochondrial genomes, with emerging research showing that they may be common in eukaryotic genomes as well [10], [20], [31], [32], [37]. Studying overlapping genes can shed light on the processes of de novo gene birth [36].

Overlapping genes tend to encode proteins with higher intrinsic structural disorder (ISD) than those encoded by non-overlapping genes [36]. The term disorder applies broadly to proteins which, at least in the absence of a binding partner, lack a stable secondary and tertiary structure. There are different degrees of disorder: molten globules, partially unstructured proteins and random coils with regions of disorder spanning from short (less than 30 residues in length) to long. Disorder can be shown experimentally or predicted from amino acid sequences using software [15]. [36] estimated, using the latter approach, that 48% of amino acids in overlapping regions exhibit disorder, compared to only 23% in non-overlapping regions.

In this work we explore three non-mutually-exclusive hypotheses why this might be, and quantify the extent of each. Two have previously been considered: that elevated ISD in overlapping genes is a mechanism that relieves evolutionary constraint, and that elevated ISD is a holdover from the de novo gene birth process. We add consideration of a third, previously-unexplored hypothesis — that elevated ISD with dual-coding may be the result of an artifact of the genetic code — to the mix.

Overlapping genes are particularly evolutionarily constrained because a mutation in an overlapping region simultaneously affects both of the two (or occasionally more) genes involved in that overlap. Because ~70% of mutations that occur in the third codon position are synonymous, versus only ~5% and 0% of mutations in the first and second codon positions respectively [38], a mutation that is synonymous in one reading frame is highly likely to be nonsynonymous in another, so to permit adaptation, overlapping genes must be relatively tolerant of nonsynonymous changes. Demonstrating the higher constraint on overlapping regions, they have lower genetic diversity and *d_N_*/*d_S_* than non-overlapping regions in RNA viruses [25], [39], [40], [43].

High ISD can alleviate the problem of constraint. Amino acid substitutions that maintain disorder have a reasonable chance of being tolerated, in contrast to the relative fragility of a well-defined three-dimensional structure. This expectation is confirmed by the higher evolutionary rates observed for disordered proteins [4], [13]. The known elevation of evolutionary constraint on overlapping genes is usually invoked as the sole [47], [53] or at least dominant [36] explanation for their high ISD. Given the strength of the evidence for constraint [25], [39], [40], [43], we attribute to constraint, by a process of elimination, what cannot be explained by our other two hypotheses.

The second hypothesis that we consider is that high ISD in overlapping genes is an artifact of the process of de novo gene birth [36]. There is no plausible path by which two non-overlapping genes could re-encode an equivalent protein sequence as overlapping; instead, an overlapping pair arises either when a second gene is born de novo within an existing gene, or when the boundaries of an existing gene are extended to create overlap [38]. In the latter case of ‘‘overprinting” [7], [19], [36], the extended portion of that gene, if not the whole gene, is born de novo. One overlapping protein-coding sequence is therefore always evolutionary younger than the other; we refer to these as ‘novel” versus ‘ancestral” overlapping genes or portions of genes. Genes may eventually lose their overlap through a process of gene duplication followed by subfunctionalization [19], enriching overlapping genes for relatively young genes that have not yet been through this process. However, gene duplication may be inaccessible to many viruses (in particular, many RNA, ssDNA, and retroviruses), due to intrinsic geometric constraints on maximum nucleotide length [6], [9], [11].

Young genes are known to have higher ISD than old genes, with high ISD at the moment of gene birth facilitating the process [52], perhaps because cells tolerate them better [48]. Domains that were more recently born de novo also have higher ISD [3], [5], [14], [26]. High ISD could be helpful in itself in creating novel function, or it could be a byproduct of a hydrophilic amino acid composition whose function is simply the avoidance of harmful protein aggregation [16], [22]. Regardless of the cause of high ISD in young genes, the “facilitate birth” hypothesis makes a distinct prediction from the constraint hypothesis, namely that the novel overlapping reading frames will tend to encode higher ISD than the ancestral overlapping reading frames.

Under the constraint hypothesis, ancestral overlapping proteins will still have elevated ISD relative to non-overlapping proteins, even if it is less elevated than that of novel overlapping proteins. Elevated ISD in the ancestral member of the gene pair might have already been there at the moment of gene birth, or it might have subsequently evolved, representing two subhypotheses within the overall hypothesis of constraint. The overlapping gene pairs that we observe are those that have been retained; if either member of the overlapping pair was born with low ISD, then constraint makes it difficult to adapt to a changing environment, and that pair is less likely to be retained. When the ancestral member of the pair already has high ISD at the moment at which the novel gene is functionalized, long-term maintenance of both genes in the face of constraint is more likely. The ‘already there” or “preadaptation” [52] version of the constraint hypothesis predicts that the pre-overlapping ancestors of today’s ancestral overlapping genes had higher ISD than other genes, perhaps because these gene pairs are the ones to have been retained. While these ancestral sequences are not available, as a proxy we use homologous sequences from basal lineages whose most recent common ancestry predates the origin of the overlap. For simplicity, we refer to these sequences as “pre-overlapping” to distinguish them from “non-overlapping” genes never known to have overlap. The preadaptation version of the constraint hypothesis predicts higher ISD in pre-overlapping genes than in non-overlapping genes.

Finally, here we also consider the possibility that the high ISD observed in overlapping genes might simply be an artifact of the genetic code [21]. We perform for the first time the appropriate control, by predicting what the ISD would be if codons were read from alternative reading frames of existing non-overlapping genes. Any DNA sequence can be read in three reading frames on each of the two strands, for a total of 6 reading frames. We focus only on same-strand overlap, due to superior availability of reliable data on same-strand overlapping gene pairs. We classify the reading frame of each gene in an overlapping pair relative to its counterpart; if gene A is in the +1 frame with respect to gene B, this means that gene B is in the +2 frame with respect to gene A. We use the +0 frame designation just for nonoverlapping or pre-overlapping genes in their original frame. If the high ISD of overlapping genes is primarily driven by the intrinsic properties of the genetic code, then we expect their ISD values to closely match those expected from translation in the +1 vs. +2 frames of non-overlapping genes.

Here we test the predictions of all three hypotheses, as summarized in Figure 1, and find that both the birth-facilitation and conflict-resolution hypotheses play a role. The artifact hypothesis plays no appreciable role in elevating the ISD of overlapping regions; while reading frame (+1 vs. +2) strongly affects the ISD of individual genes, each overlapping gene pair has one of each, and the two cancel out such that there is no net contribution to the high ISD found in overlapping regions. Surprisingly, novel genes are more likely to be born in the frame prone to lower ISD; this seems to be a case where mutation bias in the availability of open reading frames (ORFs) is more important than selection favoring higher ISD for novel than ancestral genes.

**Fig. 1.**
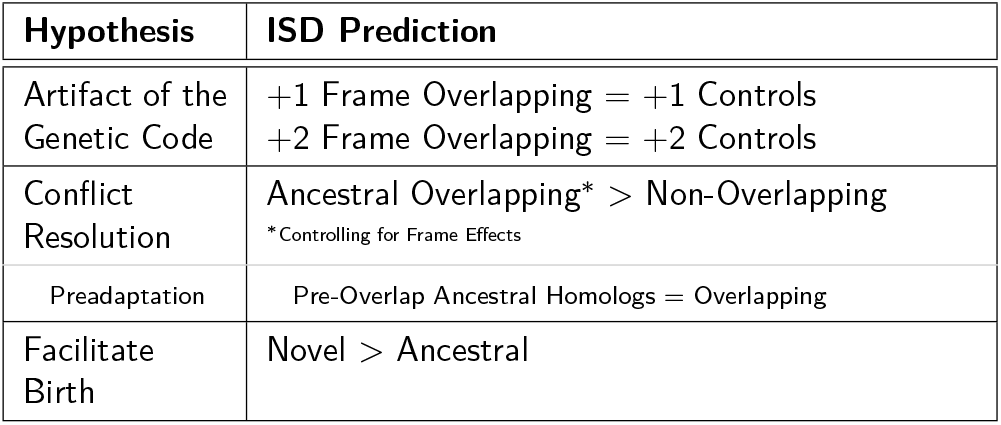
Three non-mutually-exclusive hypotheses about why overlapping genes have high ISD. The column on the right describes the ISD patterns we would expect if the hypotheses were true.

## II. Materials and Methods

### A. Overlapping Viral Genes

A total of 102 viral same-strand overlapping gene pairs were compiled from the literature [35], [36], [40], [42], [43], [51]. Of these, ten were discarded because one or both of the genes involved in the overlap were not found in the ncbi databases, either because the accession number had been removed, or because the listed gene could not be located. This left 92 gene pairs for analysis from 80 different species, spanning 33 viral families. Six of these pairs were ssDNA, five were retroviruses, while the remaining 81 were RNA viruses: 7 dsRNA, 61 positive sense RNA and 13 negative sense RNA.

### B. Relative Gene Age

For 39 of the remaining 92 gene pairs available for analysis, the identity of the older vs. younger member of the pair had been classified in the literature [27], [36], [40], [42] via phylogenetic analysis. There was disagreement in the literature regarding the TGBp2/TGBp3 overlap; we followed [27] rather than [36].

We also used relative levels of codon bias to classify the relative ages of members of each pair. Because all of the overlapping genes are from viral genomes, we can assume that they are highly expressed, leading to a strong expectation of codon bias in general. Novel genes are expected to have less extreme codon bias than ancestral genes due to evolutionary inertia [35], [40].

For each viral species, codon usage data [29], [55] were used to calculate a relative synonymous codon usage (RSCU) value for each codon [17]:

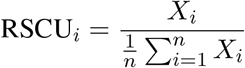

where *X_i_* is the number of occurrences of codon i in the viral genome, and 1 ≤ *n* ≤ 6 is the number of synonymous codons which code for the same amino acid as codon *i*. The relative adaptedness value (*w_i_*) for each codon in a viral species was then calculated as:

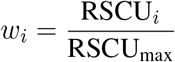

where RSCU_max_ is the RSCU value for the most frequently occurring codon corresponding to the amino acid associated with codon *i*. The codon adaptation index (CAI) was then calculated for the overlapping portion of each gene, as the geometric mean of the relative adaptedness values:

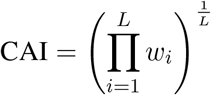

where *L* is the number of codons in the overlapping portion of the gene, excluding ATG and TGG codons. This exclusion is because ATG and TGG are the only codons that code for their respective amino acids and so their relative adaptedness values are always 1, thereby introducing no new information. To ensure sufficient statistical power to differentiate between CAI values, we did not analyze CAI for gene pairs with overlapping sections less than 200 nucleotides long.

Within each overlapping pair, we provisionally classified the gene with the higher CAI value as ancestral and the gene with lower CAI value as novel. We then compared the two sets of relative adaptedness valuesusing the wilcox.test function in R to perform a Mann-Whitney U Test. We chose a p-value cutoff of 0.035 after analyzing a receiver operating characteristic (ROC) plot (Figure 2A). The combined effects of our length threshold and p-value cutoff are illustrated in Figure 2B. In total, 27 gene pairs were determined to have statistically-significant CAI values, 19 of which had also been classified via phylogenetic analysis.

**Fig. 2.**
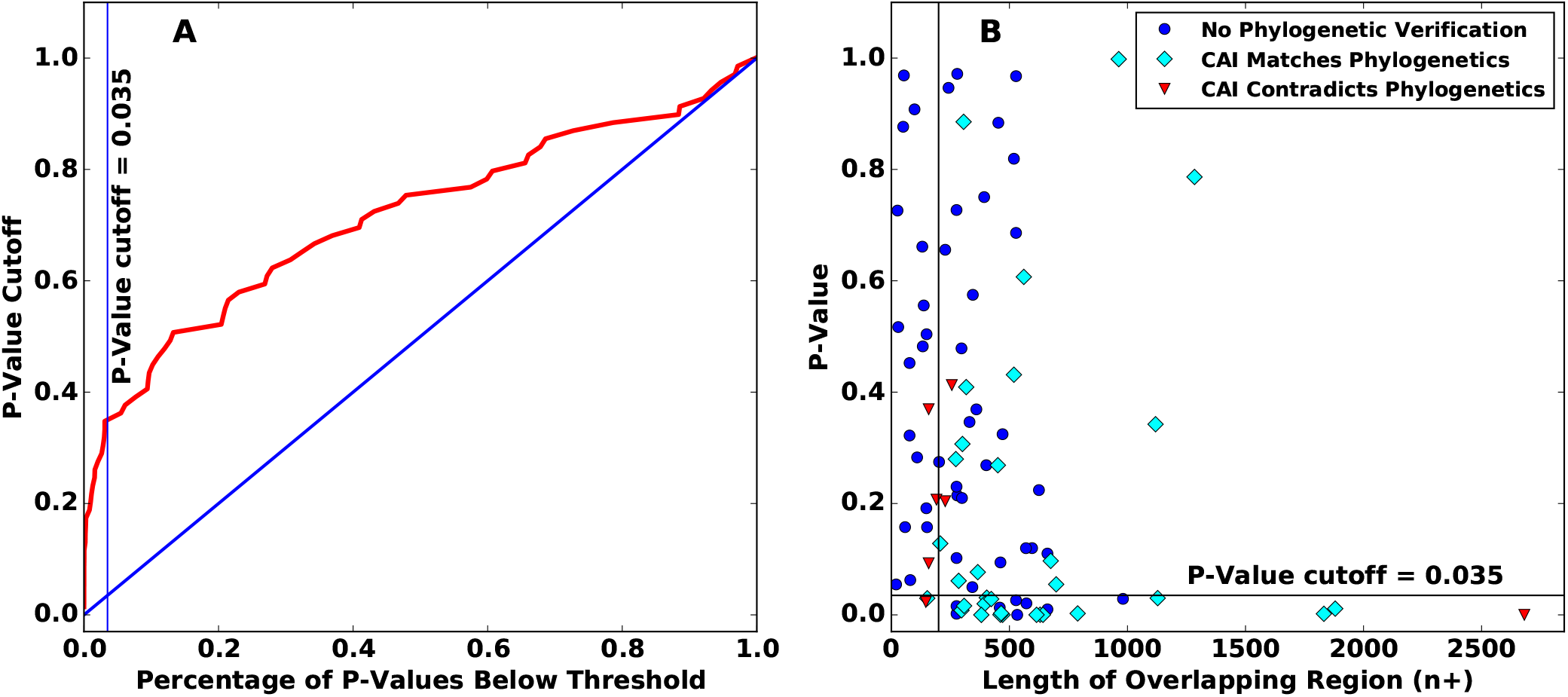
Statistical classification of relative ages. (**A**) The receiver operating characteristic plot for determining which member of an overlapping gene pair has higher CAI, and is hence presumed to be ancestral. Only genes with an overlapping region of at least 200 nucleotides are plotted. (**B**) CAI classification of the 91 gene pairs for which codon usage data were available was based both on p-value and on the length of the overlapping regions. The vertical line shows the overlapping length cutoff of 200 nucleotides, the horizontal line shows the p-value cutoff; CAI classification was considered informative for the 27 bottom right points.

Of the gene pairs whose ancestral vs. novel classification were obtained both by statistically significant CAI differences and by phylogenetics, there was one for which the CAI classification contradicted the phylogenetics. That exception was the p104/p130 overlap in the Providence virus. This overlap is notable because the ancestral member of the pair was acquired through horizontal gene transfer, which renders codon usage an unreliable predictor of relative gene ages [35]. We therefore used the phylogenetic classification and disregarded the CAI results. In total, we were able to classify ancestral vs. novel status for 47 overlapping gene pairs (Figure 3).

**Fig. 3.**
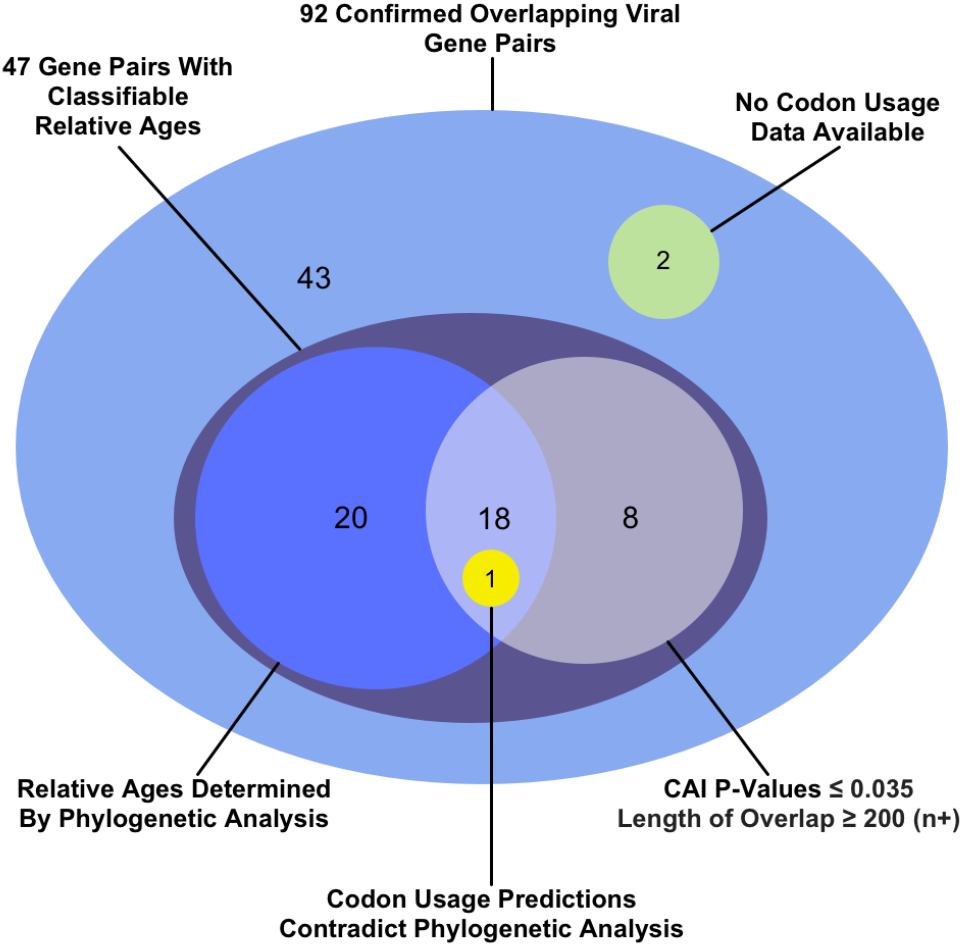
How the relative ages of the genes were classified for 47 out of the 92 overlapping gene pairs for which sequence data were available. Within each shaded region, each gene pair is counted within only one of the subregions shown. Each shaded region’s total is found by summing the individual subtotals within it, some of which are noted outside the shaded regions. For example, the relative ages of 39 genes were classified via phylogenetic analysis: 20 through phylogenetic analysis alone (blue circle), 18 supported via codon analysis (intersection of blue and white circle), and one contradicted by codon analysis (yellow circle).

### C. Non-overlapping and Pre-overlapping Controls

Both non-overlapping genes and pre-overlapping genes were used as controls. 150 non-overlapping genes were compiled from the viral species in which the 92 overlapping gene pairs were found; matching for species helps control for %GC content or other idiosyncrasies of nucleotide composition.

Of the 47 overlapping gene pairs for which we could assign relative ages, we were able to locate preoverlapping homologs for 27 of the ancestral genes in our dataset in the literature [40] and/or NCBI (BLAST search with E-value threshold = 10^−6^).

Frameshifting these control sequences was performed in two ways. First, removing one or two nucleotides immediately after the start codon and two or one nucleotides immediately before the stop codon generated +1 and +2 frameshifted non-ORF controls, respectively (Figure 4A). Any stop codons that appeared in the new reading frames were removed.

**Fig. 4.**
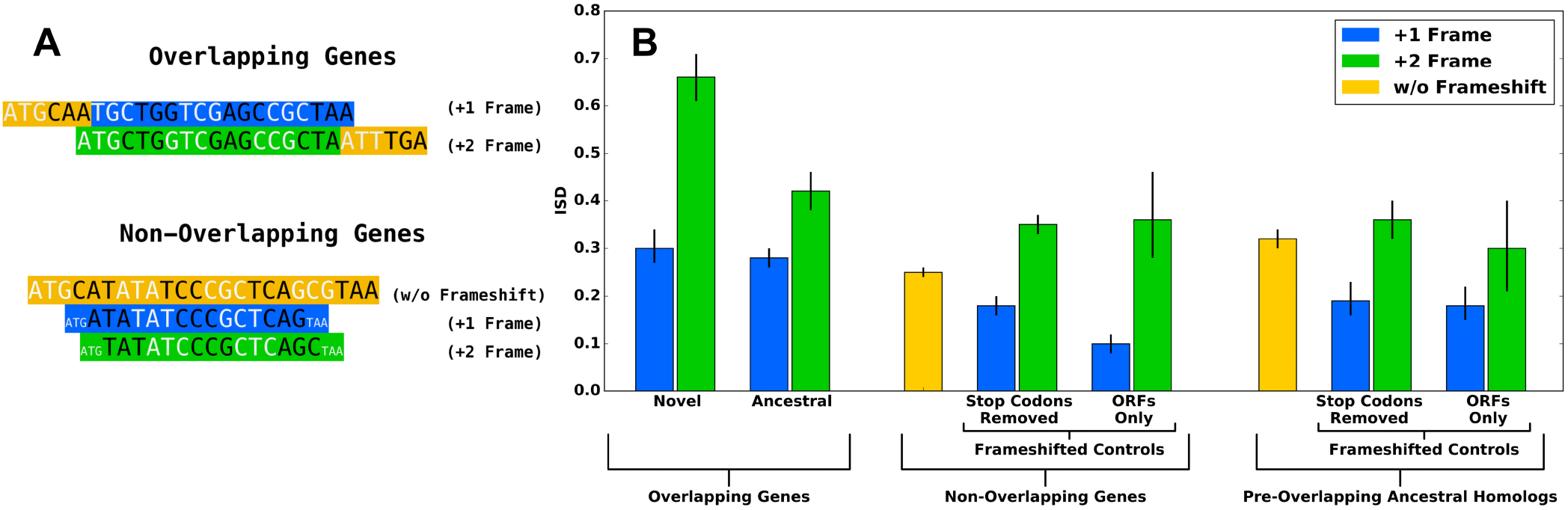
ISD results support the birth-facilitation and (preadaptation version of the) conflict resolution hypotheses. (A) Data are from the overlapping sections of the 47 gene pairs whose ages could be classified, from non-overlapping genes in the species in which those 47 overlapping gene pairs were found, and from the 27 available pre-overlapping ancestral homologs. Some numbers in the main text are for the full set of 92 overlapping gene pairs and might not match exactly. (B) While frame significantly impacts disorder content (green is consistently higher than blue), it does not drive the high ISD of overlapping genes. Even controlling for frame, novel ISD > ancestral (left), supporting birth facilitation. Supporting conflict resolution, ancestral > non-overlapping either in-frame (yellow) or matched frameshifted control. The preadaptation version of the constraint hypothesis is supported by the fact that non-overlapping ISD < pre-overlapping (compare the two yellow bars). Means and 66% confidence intervals were calculated from the Box-Cox transformed (with λ = 0.4) means and their standard errors, and are shown here following back-transformation.

Second, for the frameshifted ORF controls, we took only in situ ORFs within each of the two alternative reading frames. If multiple ORFs terminated at the same stop codon, we used only the longest. We excluded ORFs less than 25 amino acids in length (after removal of cysteines for analysis by IUPred). For nonoverlapping genes, this yielded 151 and 24 ORFs in the +1 and +2 reading frames, respectively. For the smaller set of pre-overlapping genes, it yielded only 25 and 5 ORFs in the +1 and +2 frames, respectively.

### D. Homology Groups

Treating each gene as an independent datapoint is a form of pseudoreplication, because homologous genes can share properties via a common ancestor rather than via independent evolution. This problem of phylogenetic confounding can be corrected for by using gene family as a random effect term in a linear model [52], and by counting each gene birth event only once.

We constructed a pHMMer (http://hmmer.org/) database including all overlapping regions, nonoverlapping genes and their frameshifted controls. After an all-against-all search, sequences that were identified as homologous, using an expectation value threshold of 10^−4^, were provisionally assigned the same homology group ID. These provisional groups were used to determine which gene birth events were unique. Two pairs were considered to come from the same gene birth event when both the ancestral and the overlapping sequence were classified as homologous. We also used published phylogenetic analysis to classify the TGBp2/TGBp3 overlap as two birth events (one occurring *Virgaviridae*, the other occurring in *Alpha*- and *Betaflexiviridae)* [27].

Some homologous pairs had such dissimilar protein sequences that ISD values were essentially independent. We therefore manually analyzed sequence similarity within each homology group using the Geneious [18] aligner with free end gaps, using Blosum62 as the cost matrix. The percent similarity using the Blosum62 matrix with similarity threshold 1 was then used as the criterion for whether a protein sequence would remain in its homology group for the ISD analysis. We used ≥ 50% protein sequence similarity as the threshold to assign a link between a pair, and then used single-link clustering to assign protein sequences to 561 distinct homology groups. Pre-overlapping genes were then assigned to the homology groups of the corresponding ancestral genes.

### E. ISD Prediction

We used IUPred [12] to calculate ISD values for each sequence. Following [52], before running IUPred, we excised all cysteines from each amino acid sequence, because of the uncertainty about their disulphide bond status and hence entropy [49]. Whether cysteine forms a disulphide bond depends on whether it is in an oxidizing or reducing environment. IUPred implicitly, through the selection of its training data set, assumes most cysteines are in disulphide bonds, which may or may not be accurate for our set of viral proteins. Because cysteines have large effects on ISD (in either direction) depending on disulphide status and hence introduce large inaccuracies, cysteines were dropped from consideration altogether.

IUPred assigns a score between 0 and 1 to each amino acid. To calculate the ISD of an overlapping region, IUPred was run on the complete protein (minus its cysteines), then the average score was taken across only the pertinent subset of amino acids.

### F. Statistical Models

Prior to fitting linear models, sequence-level ISD values were transformed using a Box-Cox transform. The optimal value of λ for the combined ancestral, novel and artificially-frameshifted non-overlapping, non-ORF control group data was 0.41, which we rounded to 0.4 and used throughout all linear models, and for central tendency and confidence intervals in the figures. Simple means and standard errors are described in-line in the text.

We used a multiple regression approach to determine which factors predict ISD values [44]. Gene designation (ancestral vs. novel vs. one or more non-genic controls) and relative reading frame (+1 vs. +2) were used as fixed effects. Homologous sequences are not independent; we accounted for this by using a linear mixed model [34], with homology group as a random effect. Species is a stand-in for a number of confounding factors, e.g. %GC content, and so was included as a second random effect. The data were normalized using a Box-Cox transformation prior to analysis. Pairwise comparisons discussed throughout the Results were performed using contrast statements applied to the linear model, using the minimum number of gene designations necessary to make the comparison in question.

We used the lmer and gls functions contained in the nlme and lme4 R packages to generate the linear mixed models. The main model used to calculate the relative effect sizes used frameshifted non-ORF nonoverlapping genes as the non-genic control. In this model, frame, gene designation, species, and homology group terms were retained in the model with *p* = 5×10^−20^, 1×10^−6^, 9×10^−19^, and 2×10^−3^ respectively. All four terms were also easily retained in all other model variants exploring different non-genic controls.

We also considered overlap type (internal vs. terminal) as a fixed effect, but removed it because it did not significantly enhance our model (*p* = 0.86). To determine the statistical significance of each effect, we used the anova function in R to compare nested models.

### G. Data Availability

Scripts and data tables used in this work may be accessed at: https://github.com/MaselLab/Willis_Masel_Overlapping__Genes_Structural_Disorder_Explained

## III. Results

### A. Confirming elevated ISD in our dataset

Because most verified gene overlaps in the literature, especially longer overlapping sequences, are in viruses [30], [36], [50], we focused on viral genomes, compiling a list of 92 verified overlapping gene pairs from 80 viral species. The mean predicted ISD of all overlapping regions (0.39 ± 0.02) was higher than that of the non-overlapping genes (0.25 ± 0.01), confirming previous findings that overlapping genes have elevated ISD.

### B. Frame affects ISD

We artificially frameshifted 150 non-overlapping viral genes in those 80 species, and found higher ISD in the +2 reading frame (0.35 ± 0.02) than in the +1 reading frame (0.19 ± 0.01) for non-ORF controls, and an even more extreme difference for ORF controls (0.47 ± 0.02 vs. 0.18 ± 0.01). The artifact hypothesis predicts that the +1 and +2 members of the 92 verified overlapping gene pairs will follow suit. In agreement with this, the overlapping regions of genes in the +2 reading frame had higher mean ISD (0.48 ± 0.03) than those in the +1 reading frame (0.31 ± 0.02). While this provides strong evidence that frame shapes ISD as an artifact of the genetic code, average ISD across both ways of frameshifting nonoverlapping genes (0.27±0.01 and 0.22 ±0.02 for non-ORF and ORF frameshifted non-overlapping genes, respectively) is significantly lower than the ISD of all overlapping sequences (0.39 ± 0.02), showing that the artifact hypothesis cannot fully explain elevated ISD in the latter.

#### The Birth-Facilitation Hypothesis is Supported

We find stronger support for the birth-facilitation hypothesis. Of the 92 verified overlapping viral gene pairs, we were able to classify the relative ages of the component genes as ancestral vs. novel for 47 pairs (Table I). In agreement with the predictions of the birth-facilitation process, and controlling for frame, novel genes have higher ISD than either ancestral members of the same gene pairs or artificially-frameshifted controls (Figure 4B). We confirmed this using a linear mixed model, with frame (+1 vs. +2) as a fixed effect, gene type (novel vs. ancestral) as a fixed effect, species (to control for %GC content and other subtle sequence biases) as a random effect, and homology group (to control for phylogenetic confounding) as a random effect. Within this linear model, the prediction unique to the birth-facilitation hypothesis, namely that ISD in the overlapping regions of novel genes is higher than that in ancestral genes, is supported with *p* = 0.03. Inspection of Figure 4B suggests that elevation of novel gene ISD above ancestral is specific to the +2 frame; this is confirmed in the analysis of Figure 5B. Running separate statistical models for the two frames, the +2 frame difference is supported with *p* = 0.006.

**TABLE I.**
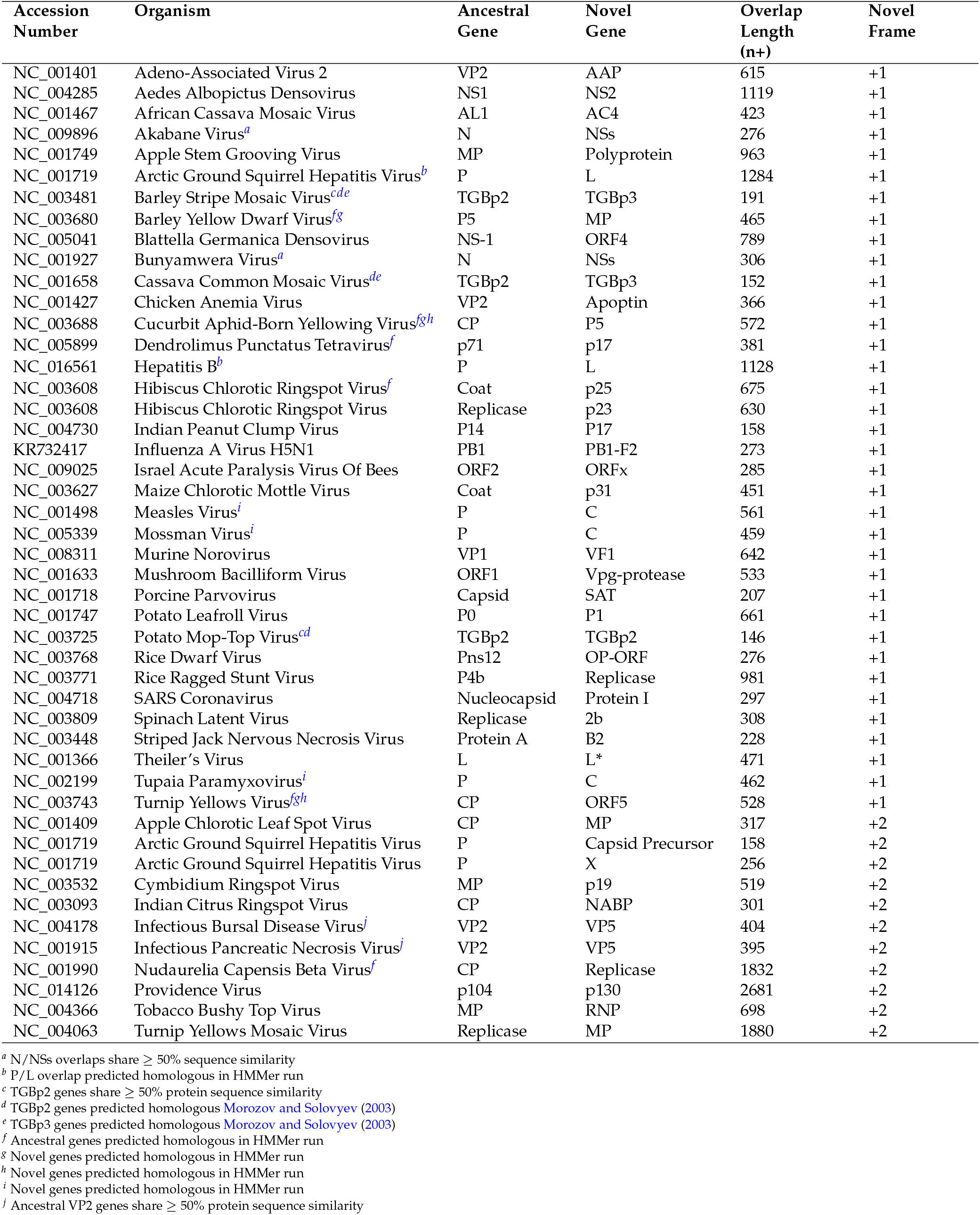
Gene pairs for which the relative ages could be determined. Genes are phylogenetically independent except as noted in the footnotes.

**Fig. 5.**
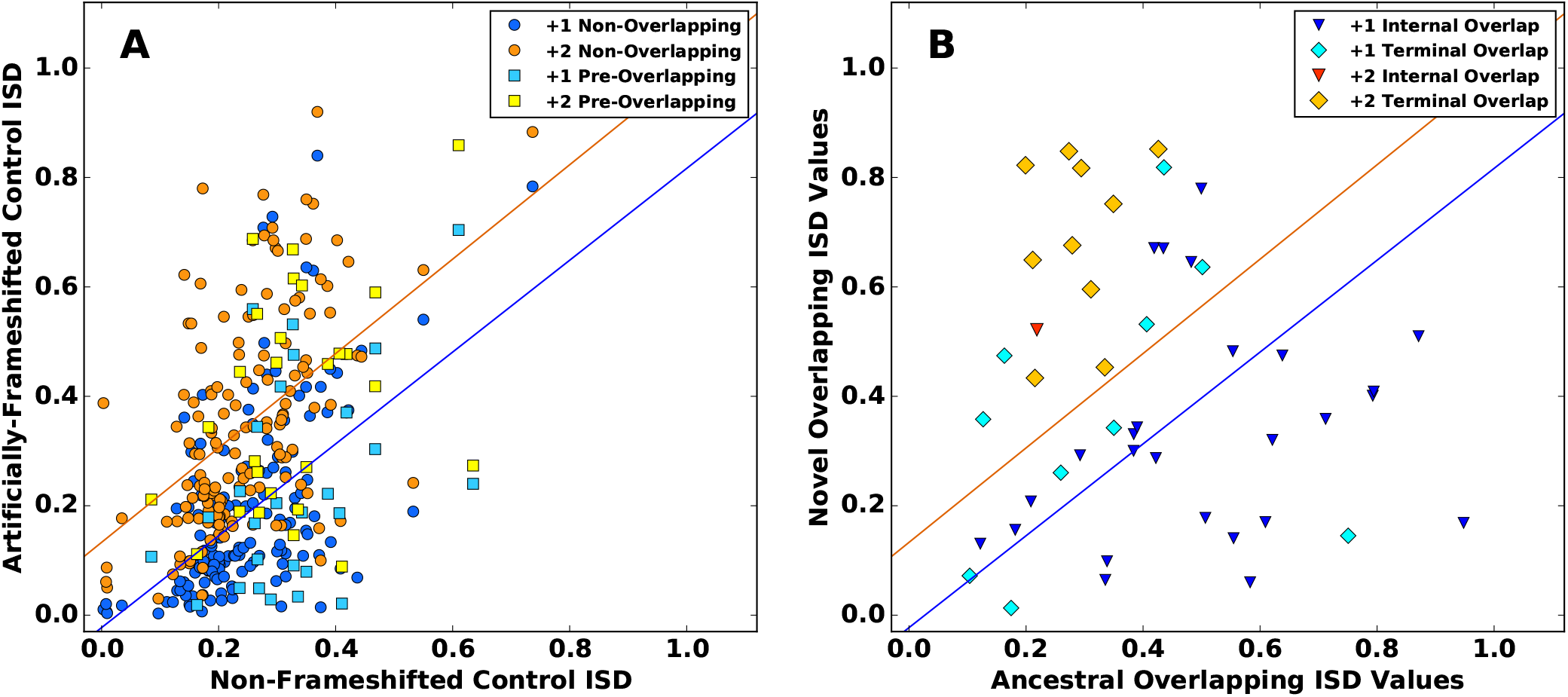
(**A**) The ISD of a non-overlapping gene predicts the ISD of the non-ORF frameshifted version of that sequence. Model I regression lines are statistically indistiguishable in our set of 150 nonoverlapping viral genes vs. our set of 27 preoverlapping genes, which are therefore pooled (*R*^2^ = 0.30 and *R*^2^ = 0.24 for the +1 and +2 groups, respectively). (**B**) The overlapping ISD of the 47 gene pairs with classifiable relative ages. Each datapoint represents one overlapping pair. The regression lines from (A) are superimposed to illustrate the elevation of novel gene ISD born into the already intrinsically high-ISD +2 frame, which destroys the correlation (*R*^2^ = 6.6 × 10^−5^ for +2 in contrast to 0.26 for +1).

These non-ORF frameshifted control sequences do not take into account the fact that ORFs vary in their propensity to appear and disappear, and that this can shape the material available for de novo gene birth, including ISD values [33]. However, we did not observe this affecting ISD in our data set. In a linear model comparing ORF vs non-ORF non-overlapping control sequences, and one comparing ORF vs. non-ORF pre-overlapping controls sequences, there was no significant difference between the two controls (*p* = 0.7 in both cases, with frame included as a fixed effect, and gene as a random effect). This justifies a focus on the larger non-ORF control dataset, as well as excluding this non-adaptive force as a driver of the birth facilitation hypothesis.

#### The Preadaptation Version of the Conflict-Resolution Hypothesis is Supported

In agreement with the conflict resolution hypothesis, ancestral overlapping sequences, not just novel ones, have higher ISD than non-overlapping genes (0.35 ± 0.02 vs. 0.25 ± 0.01; *p* = 2 × 10^−5^). This seems to be because ISD was already high at the time of de novo gene birth; today’s ancestral overlapping genes have indistiguishable ISD from their pre-overlapping homologs (*p* = 0.6), while those pre-overlapping homologs have higher ISD than non-overlapping genes (p= 2 × 10^−3^) (Figure 4B).

It is possible that the overlapping gene pairs for which we are able to identify pre-overlapping homologs are significantly younger than other overlapping gene pairs, and that there might therefore simply not have been enough time for the ancestral members of the pair to evolve higher ISD. Contradicting this, these 27 ancestral overlapping genes have indistinguishable ISD from the other 20 ancestral overlapping gene (p= 0.2, controlling for frame and homology group).

#### The Artifact Hypothesis is Rejected

Non-overlapping gene ISD is not statistically different from the mean of the +1 and +2 artificially-frameshifted control versions of the same nonoverlapping nucleotide sequences (*p* = 0.88; contrast statement applied to a linear model with fixed effect of actual non-overlapping gene sequence vs. +1 artificially-frameshifted version vs +2 artificially-frameshifted version, and gene identity as a random effect). In other words, despite the enormous effect of +1 vs. +2 reading frame, we find no support for the artifact hypothesis in explaining the elevated ISD of overlapping regions. In each overlapping gene pair, there is always exactly one gene in each of the two reading frames, such that the large effects of each of the two frames cancel each other out when all overlapping genes are considered together. It is nevertheless important to control for the large effect of frame while testing and quantifying other hypotheses.

#### The Relative Magnitude of Each Hypothesis

We calculated the degree to which birth facilitation elevates ISD using a contrast statement, as half the difference between novel and ancestral genes, because exactly half of the genes are novel, and hence elevated above the “normal” ISD level of ancestral genes. By this calculation, birth facilitation accounts for 32% ± 13% of the estimated total difference in ISD between overlapping and non-overlapping genes.

Note that frameshifted versions of high-ISD proteins have higher ISD than frameshifted versions of low-ISD proteins (Figure 5A). The two members of an overlapping pair share the same %GC content, and random sequences with higher %GC have substantially higher ISD [1], so this could be responsible for the trait correlation. A facilitate-birth bias toward high ISD in newborn genes might do so in part via high %GC in the overlapping region at the time of birth, causing overlapping sequences to be biased not just toward high-ISD novel genes, but also toward high-ISD ancestral genes. Our 32% estimate attributes all of the ISD elevation in ancestral overlapping genes to constraint, but given the trait correlation shown in Figure 5A, some of this might also be due to birth facilitation, making 32% an underestimate. Note that novel genes born into the +2 frame have high ISD above and beyond the intrinsic correlation (Figure 5B), and are thus responsible for the statistical difference between ancestral vs. novel when frame is controlled for (Figure 4B).

#### C. Mutation Bias is Responsible for More Births in the Low-ISD +1 Frame

Given the strong influence of frame combined with support for the facilitate-birth hypothesis, we hypothesized that novel genes would be born more often into the +2 frame (Figure 4A, green) because the intrinsically higher ISD of the +2 reading frame would facilitate high ISD in the novel gene and hence birth. Our dataset contained 41 phylogenetically independent overlapping pairs. Surprisingly, we found the opposite of our prediction: 31 of the novel genes were in the +1 frame of their ancestral counterparts, while only 10 were in the +2 frame (*p* = 10^−3^, cumulative binomial distribution with trial success probability 0.5).

This unexpected result is stronger for “internal overlaps”, in which one gene is completely contained within its overlapping partner (23 +1 events vs. 1 +2 event, p = 3 × 10^−6^), and is not found for ‘‘terminal overlaps”, in which the 5’ end of the downstream gene overlaps with the 3’ end of the upstream member of the pair (9 +1 events vs. 9 +2 events). (This double-counts a +1 event for which there were three homologous gene pairs, two of which were internal overlaps, and one of which was a terminal overlap.) Following [2], we interpret the restriction of this finding to internal overlaps as evidence that the cause of the bias applies to complete de novo gene birth, but not to the addition of a sequence to an existing gene.

The unexpected prevalence of +1 gene births, despite birth facilitation favoring +2, can be explained by mutation bias. One artifact of the genetic code is that +1 frameshifts yield more start codons and fewer stop codons, and hence fewer and shorter ORFs [2]. In our control set of 150 non-overlapping viral genes, we confirm that stop codons are more prevalent in the +2 frame (1 per 11 codons) than the +1 frame (1 per 14), decreasing the mean ORF length, and that start codons are more prevalent in the +1 frame (1 per 27 codons) than the +2 frame (1 per 111). Similar results were found in the pre-overlapping ancestral homologs, with more start codons in the +1 frame (1 per 33) than the +2 frame (1 per 169), and fewer stop codons in the +1 frame (1 per 20) than the +2 frame (1 per 13).

This is reflected in the relative numbers and sizes of our frameshifted ORF controls. Prior to implementing a minimum length requirement (see Materials and Methods), we found 465 ORFs in the +1 frame of our non-overlapping genes, with a mean and maximum length of 24 and 149 amino acids, respectively, while only 92 ORFs were found in the +2 frame with a mean and maximum length of 19 and 92 amino acids. The same pattern was found in the pre-overlapping ancestral homologs, with 65 ORFs found in the +1 frame with a mean and maximum length of 36 and 315 amino acids, vs. 13 ORFs in the +2 frame with a mean and maximum length of 36 and 173 amino acids.

## IV. Discussion

There is growing interest in the topic of de novo gene birth, but identifying de novo genes is plagued with high rates of both false positives and false negatives [24], with phylostratigraphy tools being particularly controversial due to homology detection biases [28]. The overlapping viral genes that we study are unlikely either to be non-genes, and must have arisen via de novo gene birth, and so circumvent many of these difficulties. [8] have disputed that young genes have high ISD, in an analysis that was prone to false positives [52]; our findings here provide an independent line of evidence, free from the danger of homology detection bias, that younger genes have higher ISD.

The study of overlapping genes has of course its own statistical traps. In particular, the preponderance of novel genes in the +1 frame demonstrates the need to control for the strong effects of frame when testing hypotheses. Ancestral genes are more frequently in the high-ISD +2 frame, while the depressed ISD of the +1 frame lowers the ISD of the novel. As a result, when frame is not considered, ancestral and novel overlapping sequences encode very similar levels of disorder (0.41 ± 0.03 vs. 0.42 ± 0.04, respectively), making it easy to miss the evidence for the facilitate-birth hypothesis.

More broadly, our results are consistent with a major role for mutational availability in shaping adaptive evolution. Rare adaptive changes happen at a rate given by the product of mutation and the probability of fixation, with the latter proportional to the selection coefficient [23]. This means that differences in the beneficial mutation rate are just as important as differences in the selection coefficient in determining which path adaptive evolution takes [54]. The influence of mutational bias has previously been observed for beneficial mutations to single amino acids in the laboratory [41], [46] and in the wild [45]. Here we demonstrate it for more radical mutations, namely the de novo birth of entire protein-coding genes.

